# Large-scale comparative genomics of *Salmonella enterica* to refine the organization of the global *Salmonella* population structure

**DOI:** 10.1101/2021.09.30.462489

**Authors:** Chao Chun Liu, William W.L. Hsiao

## Abstract

1.

Since the introduction of the White-Kauffmann-Le Minor (WKL) scheme for *Salmonella* serotyping, the nomenclature remains the most widely used for reporting the disease prevalence of *Salmonella enterica* across the globe. With the advent of whole genome sequencing (WGS), traditional serotyping has been increasingly replaced by in-silico methods that couple the detection of genetic variations in antigenic determinants with sequence-based typing. However, despite the integration of genomic-based typing by in-silico serotyping tools such as SeqSero2 and SISTR, in-silico serotyping in certain contexts remains ambiguous and insufficiently informative due to polyphyletic serovars. Furthermore, in spite of the widespread acknowledgement of polyphyly from genomic studies, the serotyping nomenclature remains unaltered. To prompt refinements to the *Salmonella* typing nomenclature for disease reporting, we herein performed a systematic characterization of putative polyphyletic serovars and the global *Salmonella* population structure by comparing 180,098 *Salmonella* genomes (representing 723 predicted serovars) from GenomeTrakr and PubMLST databases. We identified a range of core genome MLST typing thresholds that result in stable population structure, potentially suitable as the foundation of a genomic-based typing nomenclature for longitudinal surveillance. From the genomic comparisons of hundreds of predicted serovars, we demonstrated that in-silico serotyping classifications do not consistently reflect the population divergence observed at the genomic level. The organization of *Salmonella* subpopulations based on antigenic determinants can be confounded by homologous recombination and niche adaptation, resulting in shared classification of highly divergent genomes and misleading distinction between highly similar genomes. In consideration of the pivotal role of *Salmonella* serotyping, a compendium of putative polyphyletic serovars was compiled and made publicly available to provide additional context for future interpretations of in-silico serotyping results in disease surveillance settings. To refine the typing nomenclatures used in *Salmonella* surveillance reports, we foresee an improved typing scheme to be a hybrid that integrates both genomic and antigenic information such that the resolution from WGS is leveraged to improve the precision of subpopulation classifications while preserving the common names defined by the WKL scheme. Lastly, we stress the importance of controlled vocabulary integration for typing information in open data settings in order for the global *Salmonella* population dynamics to be fully trackable.

**Impact Statement:** *Salmonella enterica* (*S. enterica*) is a major foodborne pathogen responsible for an annual incidence rate of more than 90 million cases of foodborne illnesses worldwide. To surveil the high order *Salmonella* lineages, compare disease prevalence across jurisdictions worldwide, and inform risk assessments, in-silico serotyping has been established as the gold standard for typing the bacteria. However, despite previous *Salmonella* genomic studies reporting discordance between phylogenomic clades and serovars, refinements have yet been made to the serotyping scheme. Here, we analyzed over 180,000 *Salmonella* genomes representing 723 predicted serovars to subdivide the population into evolutionarily stable clusters in order to propose a stable organization of the *Salmonella* population structure that can form the basis of a genomic-based typing scheme for the pathogen. We described numerous instances in which genomes between serotypes are more similar than genomes within a serotype to reflect the inconsistencies of subpopulation classifications based on antigenic determinants. Moreover, we found inconsistencies between predicted serovars and reported serovars which highlighted potential errors in existing in-silico serotyping tools and the need to implement controlled vocabularies for reporting *Salmonella* subtypes in public databases. The findings of our study aim to motivate the future development of a standardized genomic-based typing nomenclature that more accurately captures the natural populations of *S. enterica*.

**Data Summary:** The assembly accession numbers of the genomes analyzed in this study (n = 204,952) and the associated metadata (e.g. sampling location, collection date, FTP address for retrieval) are documented in Table S1. The GenomeTrakr genomes were retrieved from the National Center for Biological Information GenBank database. The PubMLST genomes were retrieved using the BIGSdb API.

## 4. Introduction

*Salmonella enterica* (*S. enterica*) is a significant public health burden that causes more than 90 million cases of foodborne infections globally every year (1). Existing surveillance networks rely on globally standardized typing nomenclatures to rapidly share information between laboratories, veterinarians and public health authorities for risk assessments and disease tracking. Since its introduction more than 80 years ago, serotyping has been the predominant method for reporting *Salmonella* disease prevalence and represents a high-order classification to subdivide the *Salmonella* population structure into characteristic groups that differ in ecological and clinical features (2). To date, the White-Kauffmann-Le Minor (WKL) scheme subdivides *S. enterica* into over 2,600 serovars based on variations in the O (somatic) and H (flagellar) antigens (3). Although *Salmonella* serotyping has been established as a central nomenclature that is useful for disease reporting and risk assessments, its inconsistencies in the representation of natural microbial populations due to the existence of polyphyletic serovars have challenged its role as the gold standard scheme for longitudinal surveillance (4–5).

Advances in DNA sequencing over the years have opened up new avenues for microbial typing. Inference of sequence types based on genetic variations in 6-10 housekeeping genes known as multi-locus sequence typing (MLST), has shown high concordance with *Salmonella* serotyping, while achieving greater discriminatory power (5). Highly clonal serovars such as Enteritidis are widely associated with a specific sequence type, while genetically diverse serovars such as Newport and Typhimurium are subdivided into multiple sequence types. Moreover, MLST represents a more robust scheme for natural population characterization compared to antigenic variations, as it overcomes the occasional failure of the serotyping scheme to demonstrate genetic similarity amongst isolates of the same serovar (5–8). In an attempt to develop a novel *Salmonella* typing scheme that better reflects real and stable subpopulations, Achtman et al. described the concept of eBurst Groups (eBGs) that represent single-linkage chains of sequence types differing by a single locus (5). A total of 138 eBGs was defined and demonstrated effective segregation of polyphyletic serovars into distinguished groups (5). Although eBG designations have been validated across >100,000 *Salmonella* genomes and determined to be an evolutionarily stable typing scheme (8), its discriminatory power remains limited, as the scheme only accounts for genetic variations in seven housekeeping genes.

Since then, increasing adoption of whole genome sequencing (WGS) for microbial characterization accompanied by the development of efficient bioinformatic tools renders strain differentiation based on a greater number of genetic markers feasible. Leveraging the plethora of genetic information made available by WGS, core genome MLST (cgMLST) extends upon traditional MLST by the inclusion of 300-3,000 loci present in 95-99% of *Salmonella* isolates in its scheme (9). And given the conservative nature of the core genome composition, it represents a stable genomic segment that can be used to infer genealogy (9). Due to its discriminatory power superior to that of classical fingerprinting methods such as pulsed-field gel electrophoresis, phage typing or multi-locus variable-number tandem repeat analysis, cgMLST has been shown to be more effective in the differentiation of closely related strains rendering it a compelling complement for outbreak investigations (10, 11). In addition, due to the aforementioned correlation between MLST and antigenic variations, cgMLST is leveraged by in-silico serotyping workflows such as SISTR (12) and presents valuable information when putative serovars cannot be discriminated based on antigenic determinants alone.

However, the application of cgMLST for the routine tracking of high order *Salmonella* lineages has not been widely explored, as the serotyping scheme has long been used for diagnostics and longitudinal surveillance. To propose an alternative genomic-based approach to assign high order *Salmonella* groupings, we characterized stable *Salmonella* subpopulations from a diverse dataset of 180,098 *Salmonella* genomes retrieved from public databases, namely GenomeTrakr and PubMLST. We applied a recently proposed statistic called neighbourhood Adjusted Wallace Coefficient (nAWC) that quantifies single-linkage cluster concordance between two neighbouring distance thresholds ( *T* vs. *T* + 1) in which signatures of cluster stability would arise from the maintenance of high cluster concordance across multiple successive thresholds (13). The identification of long-range cgMLST typing thresholds that consistently yield high cluster concordance would inform an ideal cutoff to define a stable subdivision of the *Salmonella* genomes into characteristic groups. Such subdivision would consequently allow the development of a genomic-based typing scheme suitable for longitudinal surveillance.

It remains challenging to adopt a genomic-based typing scheme for longitudinal surveillance given how integral serotyping is for the current state of global surveillance, and decades of research on the epidemiological significance associated with different serovars rendering serotype assignments irreplaceable. Although existing in-silico serotyping tools leverage the correlation between genome-wide variations and serovars for in-silico prediction, this observed correlation has yet been systematically evaluated across most serotypes on a large scale. Hence, to contribute to the effort, we compared genomic clusters to in-silico serovar assignments in order to assess whether isolates of different serovars converge in the same genomic cluster and the extent of sequence dissimilarity amongst isolates of the same serovar. We further demonstrated the utility of existing genome-wide association methods to characterize genetic features associated with microbial lineage divergences to formulate the basis of lineage-specific subtyping schemes. The usage of typing nomenclatures that consistently represent natural bacterial populations will permit a more precise categorization of genomes and avoid the misinterpretation of epidemiological patterns from surveillance data. However, the benefits of genomic-based typing schemes cannot be realized without the community convergence on an universal genomic-based typing scheme and collaborative efforts to characterize the functional differences, ecological adaptations and health risks associated with distinct subpopulations.

## 5. Methods

### 5.1. Study dataset

The metadata of the *Salmonella* genomes archived in the GenomeTrakr database (n = 186,312) was retrieved from the National Center for Biotechnology Information (NCBI) Pathogen Detection portal (https://www.ncbi.nlm.nih.gov/pathogens) on Dec 10th, 2019. The metadata of the *Salmonella* genomes archived in the PubMLST database (n = 18,640) were retrieved from the PubMLST database (https://pubmlst.org/) on Nov 9th, 2019. The GenomeTrakr genomes were downloaded from the NCBI GenBank database on December 13th, 2019. The PubMLST genomes were retrieved using the BIGSdb API on December 13th, 2019. The associated metadata of the combined list of genomes (n = 204,952) analyzed in this study is available in Table S1. Assembly statistics such as sequence length and N50 were evaluated using QUAST v5.0.2 (14). Genome completeness and contamination were evaluated using CheckM v1.1.2 (15). The minimum data quality criteria are as follows: genome length > 4 Mbp, genome completeness > 80%, genome contamination < 5%, N50 > 50 Kbp, proportion of N bases < 1%. Assemblies that failed to meet the minimum data quality were removed prior to any downstream analysis. Following cgMLST allele calling, genomes with <5% cgMLST profile completeness were also removed from further analyses. Of the remaining 180,098 *Salmonella* genomes that passed the quality filters, the dataset is composed of 723 different serovars predicted by SISTR v1.1.1 with confidence (12). The criteria for confident SISTR prediction is determined by the presence of O/H antigenic determinants and >297/330 core genes in the query genome (12). The samples were collected from 137 countries with a sampling date range from 1900 to 2019. As the majority of contributors to the GenomeTrakr network are situated in the United States coupled with routine data synchronization with its United Kingdom’s counterpart, Enterobase, 75% of the samples were collected from either the United States or the United Kingdom (16). The samples were sequenced from 4,634 different types of isolation sources distributed across human, environment, animal, and food origins with ~60% of the samples reported as human clinical samples.

### 5.2. cgMLST and Phylogenetic analysis

The cgMLST profile of each genome was generated using chewBACCA v2.0.16 based on a 3,246 loci scheme (17). The original cgMLST scheme was downloaded from https://zenodo.org/record/1323684 and consisted of 3,255 loci; however, 18 loci appeared to be paralogous pairs and hence one loci of each paralogous pair was arbitrarily removed from the original scheme. Hamming distances were calculated to compare the allelic profile differences and generate a pairwise distance matrix. A neighbour joining tree was constructed from the pairwise distance matrix using Rapidnj v2.3.2 and visualized in GrapeTree v2.1 (18, 19). Core genome SNP phylogenomics analysis of London/Winterthur and Chester/Madras genomes was conducted using PhaME v1.0.3 which uses Mummer v3.23 to generate core genome SNP alignments from draft genomes, and FastTree v2.1.9 to construct a maximum likelihood tree (20–22).

### 5.3. Cluster stability quantification

The adjusted Wallace coefficient was initially proposed to assess cluster concordance and has since been applied to compare the results of different subtyping methods. nAWC represents an extension to the adjusted Wallace coefficient by assessing cluster concordance between neighbouring thresholds that differ by one distance unit. Cluster membership resistance to incremental changes of distance thresholds or alternatively the maintenance of high degree of cluster concordance would consequently reflect cluster stability and render the median value of the cluster stable region suitable for defining WGS-based subtyping nomenclatures for pathogen surveillance.

TreeClust was used to cluster the leaves of the constructed neighbour joining tree to generate single-linkage clusters at thresholds ranging from 1 to 3,246 allelic differences. The resultant cluster assignments were compared between neighbouring thresholds to evaluate nAWC and Shannon entropy using the R scripts available at https://github.com/theInnuendoProject/nAWC.

### 5.4. Genome-wide association and functional enrichment

PCA of the cgMLST profiles from the two Enteritidis lineages was conducted in R using the adegenet v2.1.3 package (23). The principal component loadings were analyzed to quantify the allelic contribution to the observed lineage divergence. Evaluation of genome wide association to lineage divergence was conducted using Scoary v1.6.16 that tests the statistical association between the presence or absence of genotypic features to phenotypic characteristics (24). The cgMLST profiles were transformed into an allele presence/absence matrix such that each unique allele of a loci is considered an independent genotypic feature. The Scoary output was filtered by multiple test corrected p-values to determine the set of alleles significantly associated with lineage identity. Similarly with pan-genome wide association, Roary v3.12.0 was used to generate a pan-genome gene presence/absence matrix and subsequently used as input to Scoary to identify statistical association between gene presence and lineage identity (25). The dN/dS ratio was calculated using the CODEML program part of the PAML v4.9 package (26). Protein functional annotation enrichment was performed by uploading a list of significant genes to the DAVID web service (27). COG category enrichment analysis was performed using the BOG v2.0 package in R (28).

## 6. Results

### 6.1. Genomics-based characterization of *S. enterica* subpopulations

Quantification of cluster stability by nAWC across the range of possible cgMLST distance threshold (*T*) values revealed two notable features: regions of compositionally identical clusters and cluster destabilizing thresholds (Fig. S1). Here, we define a given cluster stable region as any range of thresholds that maintains an nAWC > 0.99 for a minimum of 5 successive *T’s*. By this definition, we identified a total of 43 cluster stable regions of which the earliest signature of stability occurred at a median value of *T* = 45 allelic differences and the longest region of stability occurred at a median value of *T* = 1,538 allelic differences (Fig. S1A). The cluster membership dynamics observed with nAWC closely mirrored the changes in Shannon entropy, a commonly used statistic to quantify population structure richness and evenness. The consolidations of large clusters observed at *T* > 2,732 allelic differences demonstrated by frequent oscillations in nAWC coincided with the sharp decline of Shannon entropy (Fig. S1B). The identification of these cluster stable *T*’s provided the means to use genomics data to redefine the high order groupings of closely related *Salmonella* isolates.

Visualization of the phylogenetic tree constructed from the entire dataset revealed distinctive segregation by serovar assignments, as many serovars such as Enteritidis, Thompson, Heidelberg, Oranienburg, and Braenderup appeared to form monophyletic clades (Fig. S2). Consistent with previous reports, the frequently reported polyphyletic serovars such as Newport and Kentucky segregated into multiple lineages that differed by more than 2,000 allelic differences. Antigenically similar serovars such as variants of Paratyphi B and Typhimurium were observed to share highly similar core genomes, as the respective isolates clustered together in the tree. Another heterogeneous cluster observed in the tree is comprised of Lubbock and Mbandaka (Fig. S2) in which the observation is concordant with a previous study that suggested Lubbock have descended from a Mbandaka ancestor via a recombination event (29).

To subdivide the dataset into the most stable genomic clusters, we searched for the longest range of cgMLST threshold that maintained nAWC > 0.99, which occurred between *T* = 1,192 and *T* = 1,885. The robustness of our cluster definition derives from the fact that the cluster assignments, richness and evenness remain stable across a long range of distance cutoffs. Evaluation of the clusters defined at the median *T* of the longest region of stability (*T* = 1,538) revealed a total of 1,342 clusters and a high degree of concordance was observed between the predicted clusters and in-silico serovar assignments in which 507/723 (70.1%) serovars mapped to one specific genomics-based cluster (Fig. 1). Examples of serovars that demonstrated 1-to-1 mapping to genomic clusters included Heidelberg, Braenderup, Agona, Mbandaka and Paratyphi A. Interestingly, 15 of the top 20 most prevalent serovars that cause human salmonellosis reported by the United States Centers of Disease Control and Prevention (30) did not exhibit assignments to one specific genomic cluster at the most stable *T*, which reflected the multi-lineage nature of several clinically significant serovars, including Enteritidis, Newport, Typhimurium, i 1,4,[5],12:i:-, Javiana, Infantis, Muenchen, Montevideo, Thompson, Saintpaul, Oranienburg, Mississippi, Bareilly, Paratyphi B var. Java, and Anatum. In contrast to previous reports which often described the serovar Enteritidis to be sequence type specific and highly clonal (31), we found the Enteritidis isolates to segregate into 8 different clusters at the most stable *T;* however, significant bias was indeed observed in the relative abundance across the Enteritidis clusters in which 33,980 (99.4%) Enteritidis isolates in the dataset were classified under a single cluster, Cluster 841. It is unlikely that the minor clusters were mispredicted due to data quality, as their average genome completeness and contamination were within 1% difference from that of Cluster 841. Instead, these minor clusters could be explained by geographical origins given that the remaining 210 Enteritidis isolates were primarily sampled from Australia (40.5%), United States (27.6%), and United Kingdom (14.3%). To distinguish between uniform and skewed cluster distribution across these multi-lineage serovars, we compared the Shannon entropy of each clinically significant serovar at the most stable *T.* Of the clinically significant serovars, Bareilly, Javiana, Mississippi, Montevideo, Muenchen, Newport, Oranienburg, Paratyphi B var. Java, Saintpaul, and Thompson more closely approximated uniform cluster distribution indicating that the multi-lineage segregation of these serovars was not due to rare or undersampled distant strains (Fig. S3). Moreover, it was observed that numerous serovars were assigned to the same single-linkage clusters at the most stable *T*, as only 890/1,342 (66.3%) genomic clusters were found to consist of a single in-silico serovar. Examples included inter-serogroup clustering of serovars Bredeney (B), Give (E1), Kimuenza (B) and Schwarzengrund (B) and serogroup B clustering of serovars Brandenburg, Reading, Madras, and Sandiego.

**Figure 1.**
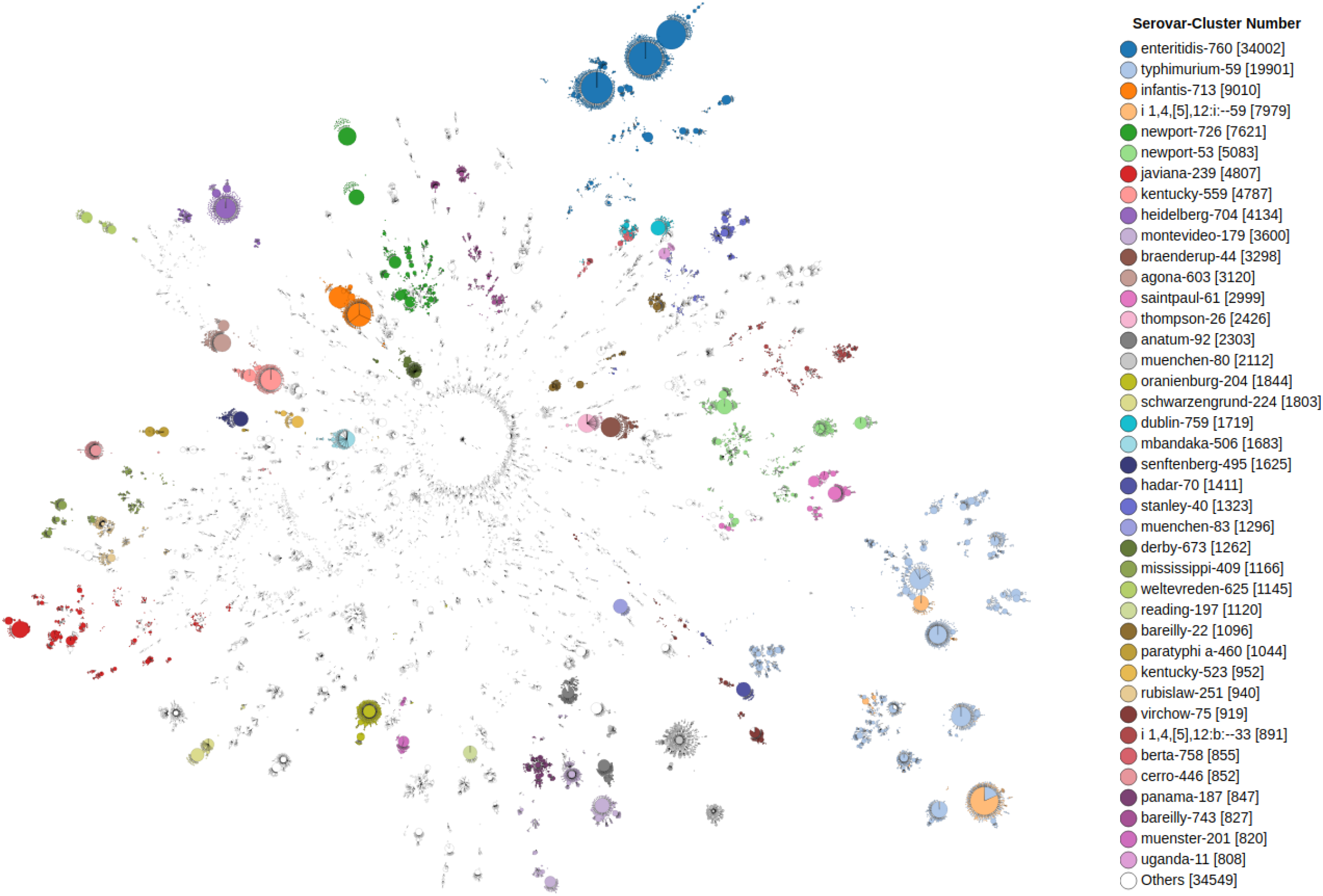
Unrooted Neighbour-joining tree constructed from 3,246-loci cgMLST profiles of 180,098 *Salmonella enterica* genomes with color annotations reflecting the combination of serovar and genomic cluster classifications. The tree is visualized using GrapeTree (43).

### 6.2. Characterization of genomically similar serovars

To comprehensively characterize a list of serovars that share highly similar core genomes, we analyzed the genomics clusters predicted at the earliest signature of stability (*T* = 45) defined by the first occurrence of 5 successive *T* with nAWC > 0.99. In this region, a total of 23,748 clusters were predicted, of which 23,584 (99.31%) clusters contained isolates of a single in-silico serovar. Subsequent investigation of the other 164 (0.69%) clusters that were compositionally mixed led to the identification of a total of 65 genomically similar serovar pairs. Of note, low confidence in-silico serovar predictions were ignored when determining compositionally mixed clusters to avoid the inference of artificial relationships due to erroneous in-silico serotyping. Monophasic/diphasic variants of Typhimurium and Paratyphi B/Paratyphi B variant Java were amongst the identified serovar pairs that shared highly similar core genomes. To evaluate the correlation between antigenic and genomic variations, the antigenic formulas of the identified genomically similar serovar pairs were compared. If the antigenic loci and genomic variations are correlated, one should expect the identified serovar pairs to share similar antigenic combinations given that the pairs are clustered based on their core genomes. Given this rationale, we identified 19 serovar pairs that differed in serogroups and 4 serovar pairs that differed by two or more antigenic types (Fig. 2). However, there were numerous genetically similar serovar pairs previously reported to share identical pulse-field gel electrophoresis patterns, but were not identified in this study. It was discovered that the insensitivity is due to consistent discrepancies between SISTR serovar predictions and the reported serovars in the metadata (Table S2). Notably, SISTR is unable to distinguish between the O-6 antigen serovar pairs reported by Mikoleit et al. (32) and consequently led to its inability to detect these genetically similar O-6 antigen serovar pairs.

**Figure 2.**
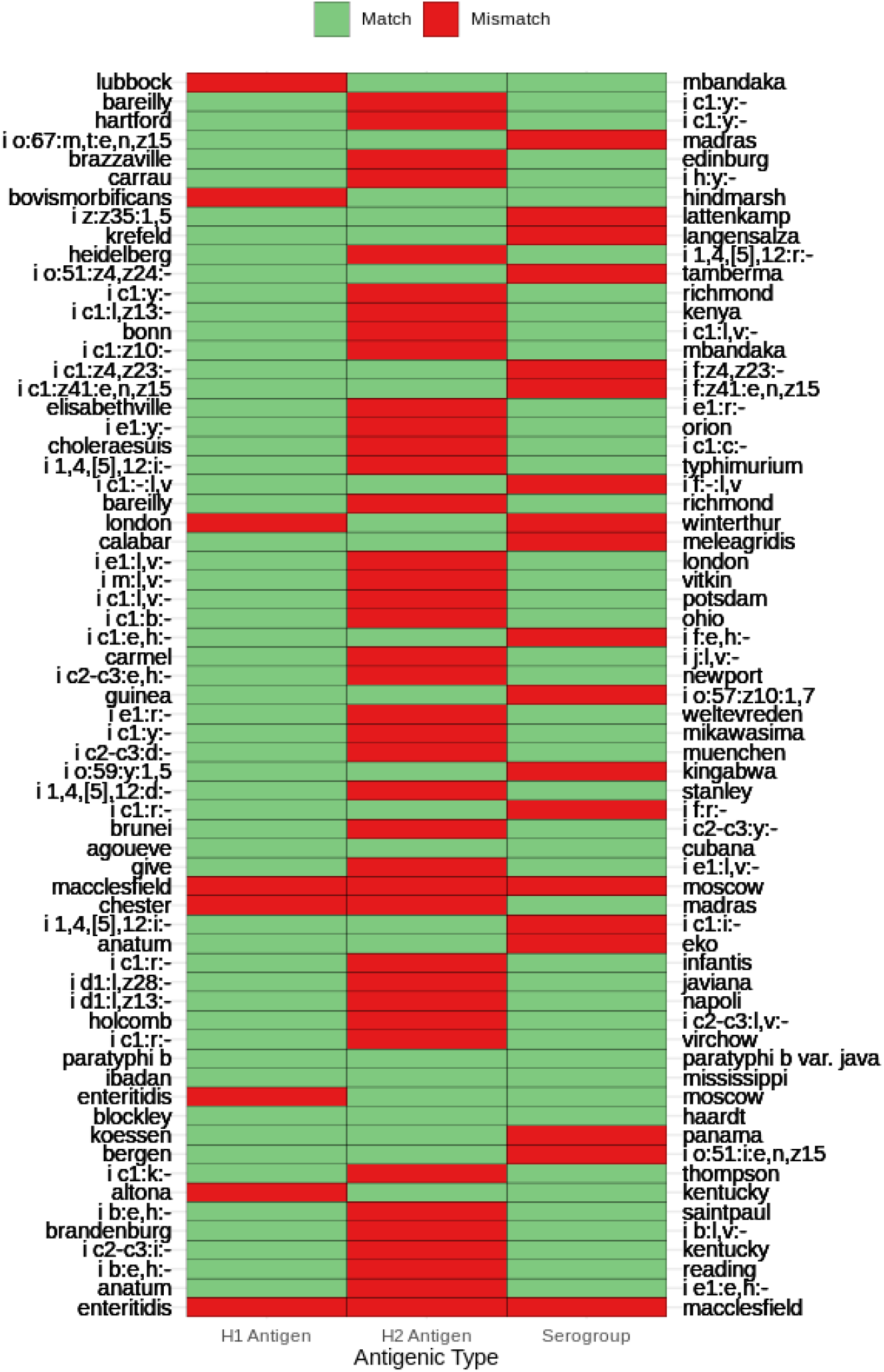
Antigenic formula comparisons of 65 genomically similar serovar pairs.

Further phylogenetic analysis of two genomically similar serovar pairs (London/Winterthur and Chester/Madras) that demonstrated unexpected clustering did not reveal evidence of a recent common descent for isolates of the same serotype. Phylogenomic analysis of all publicly available genomes for the serovar pair London (n = 412) and Winterthur (n = 2) revealed significant genetic divergence amongst serovar London and that the Winterthur isolates have likely descended from an ancestor common to a specific London lineage; however, the two Winterthur isolates were segregated and did not share a common branch in the phylogeny (Fig. S3). Similarly, the analysis of the serovar pair Chester (n = 511) and Madras (n = 6) revealed the divergence of Madras into two distant lineages of which only one appeared compositionally homogeneous (Fig. S4). Our observations of dissimilar antigens synthesized by genomically similar isolates and the inconsistent phylogenetic groupings amongst antigenically identical isolates represent examples in which predicted serovars do not represent observed phylogenetic relationships.

### 6.3. Characterization of putative non-monophyletic serovars

It is widely described in the literature that serovars have distinctive ecological and clinical features. Moreover, it has been demonstrated that serovars are genetically consistent based on MLST with a few exceptions. However, these observations have not been evaluated at a larger scale and thus may not be representative of the true genetic diversity of each serovar. Hence, to evaluate the correlation between in-silico serotyping and the lineage divergences inferred by WGS, we systematically compared the genomic similarity amongst isolates of the same in-silico serovar assignments to identify putative polyphyletic serovars and confirm whether isolates of the same serovar are in fact more similar than isolates of other serovars. Of the 738 serovars predicted by SISTR in the study dataset, 201 serovars were represented by a single sequence in the dataset and hence could not be evaluated.

We searched for putative non-monophyletic serovars by identifying occurrences in which inter-serovar comparisons yielded greater sequence identity than intra-serovar comparisons at any *T* between the earliest signature of stability ( *T* = 45) and the maximum threshold (*T* = 3,246). By this approach, we found 265/537 (49.3%) putative non-monophyletic serovars including Enteritidis, Typhimurium, Newport, Kentucky, and Javiana. Subsequent manual literature curation indicated that 18% of the evaluated serovars had been previously reported to be polyphyletic or paraphyletic (5, 33–41). Furthermore, the evaluation of the maximum distance required to link genomes of the same in-silico serovar assignments in a single cluster revealed genomes of 209/537 (38.9%) serovars required a distance of >2,000 allelic differences to be linked in a single cluster (Fig. 3). The higher than previously reported prevalence of putative polyphyletic serovars and the multitude of multi-lineage serovars demonstrated from our collective analyses express the complex nature of the evolutionary histories of *Salmonella* which cannot be comprehensively depicted by antigenic determinants. Moreover, there is also evidence to suggest that our estimation of the proportion of putative non-monophyletic serovars likely represents an underestimation of the true proportion, as serovar sample size was found to be a statistically significant predictor of non-monophyly (Fig. S5).

**Figure 3.**
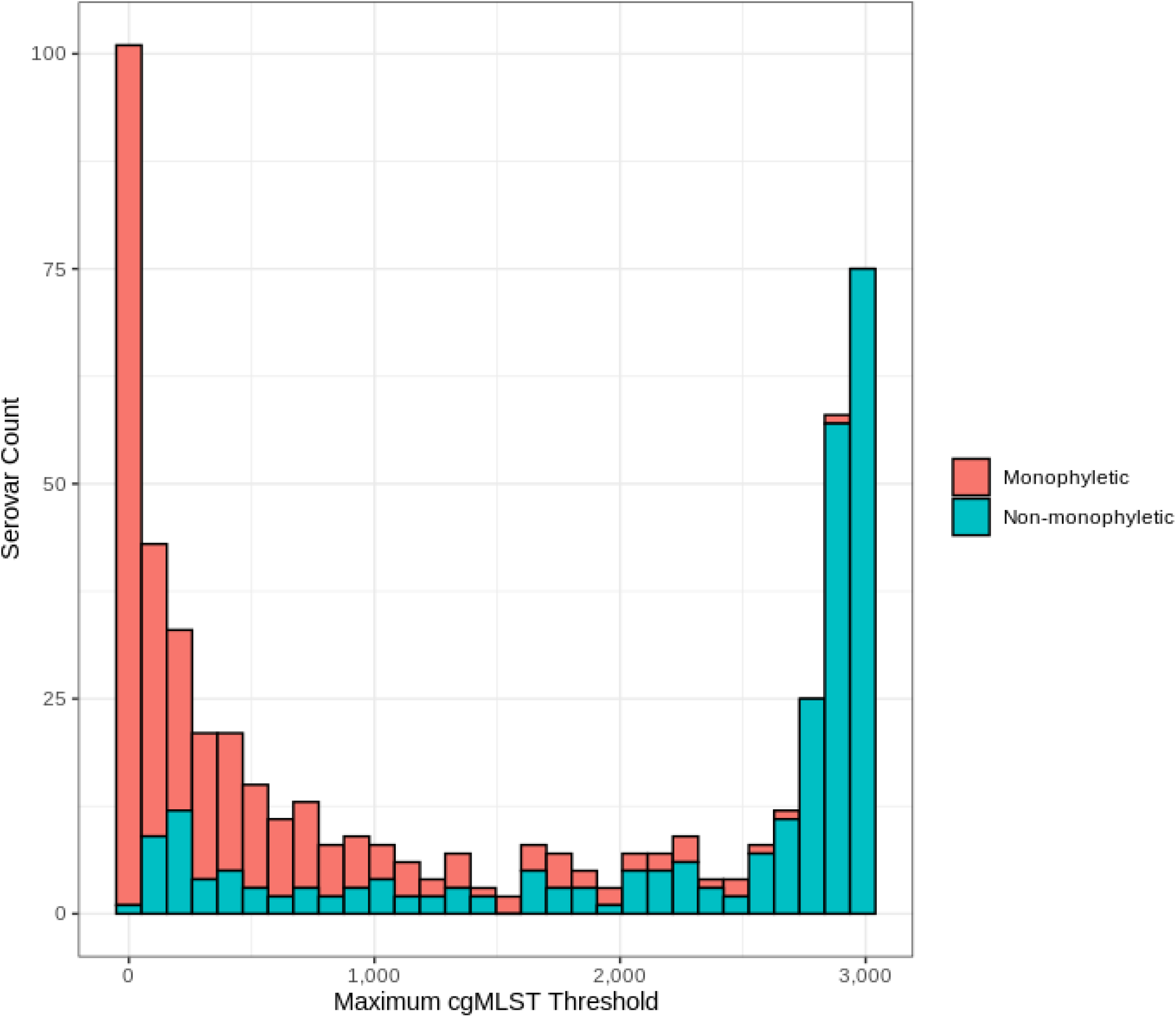
Distribution of monophyletic and non-monophyletic serovars by the maximum cgMLST allelic distance required to merge all genomes of the same serovar in a single-linkage cluster.

### 6.4. Analysis of cluster destabilizing thresholds

Significant changes in nAWC by one unit incremental change in *T* signify the consolidation of diverged lineages that differed in a set of loci. Hence, the analysis of converging clusters at cluster destabilizing thresholds can inform evolutionary-driven adaptations and divergence between diverged *Salmonella* subpopulations. Here, we analyzed a single cluster destabilizing threshold at *T* = 157 allelic differences and determined that the decrease in nAWC was due to the convergence of two Enteritidis clusters named Cluster 3139 (n = 10,546) and Cluster 3140 (n = 21,398). This was further confirmed by recomputing nAWC at *T =* 157 with the exclusion of the Enteritidis isolates from the predicted clusters which recovered an nAWC above 0.99. Comparing the geographic distribution of the two Enteritidis clusters revealed a clear geographical segregation of the two lineages of which Cluster 3139 isolates predominated in Asia (log ratio: 3.13), South America (log ratio: 3.87) and Oceania (log ratio: 4.16), while Cluster 3140 isolates predominated in North America (log ratio: 2.08) (Fig. 4A). As an orthogonal validation, principal component analysis (PCA) of the two clusters’ cgMLST profiles similarly revealed the genetic divergence of the two subpopulations that can be explained by the first principal component alone (Fig. 4B). The inclusion of the second principal component revealed the partition of Cluster 3140 into additional subpopulation structures that are suggestive of greater diversification in Cluster 3140 isolates compared to Cluster 3139 which is concordant with previous observations by Zheng et al. (42).

**Figure 4.**
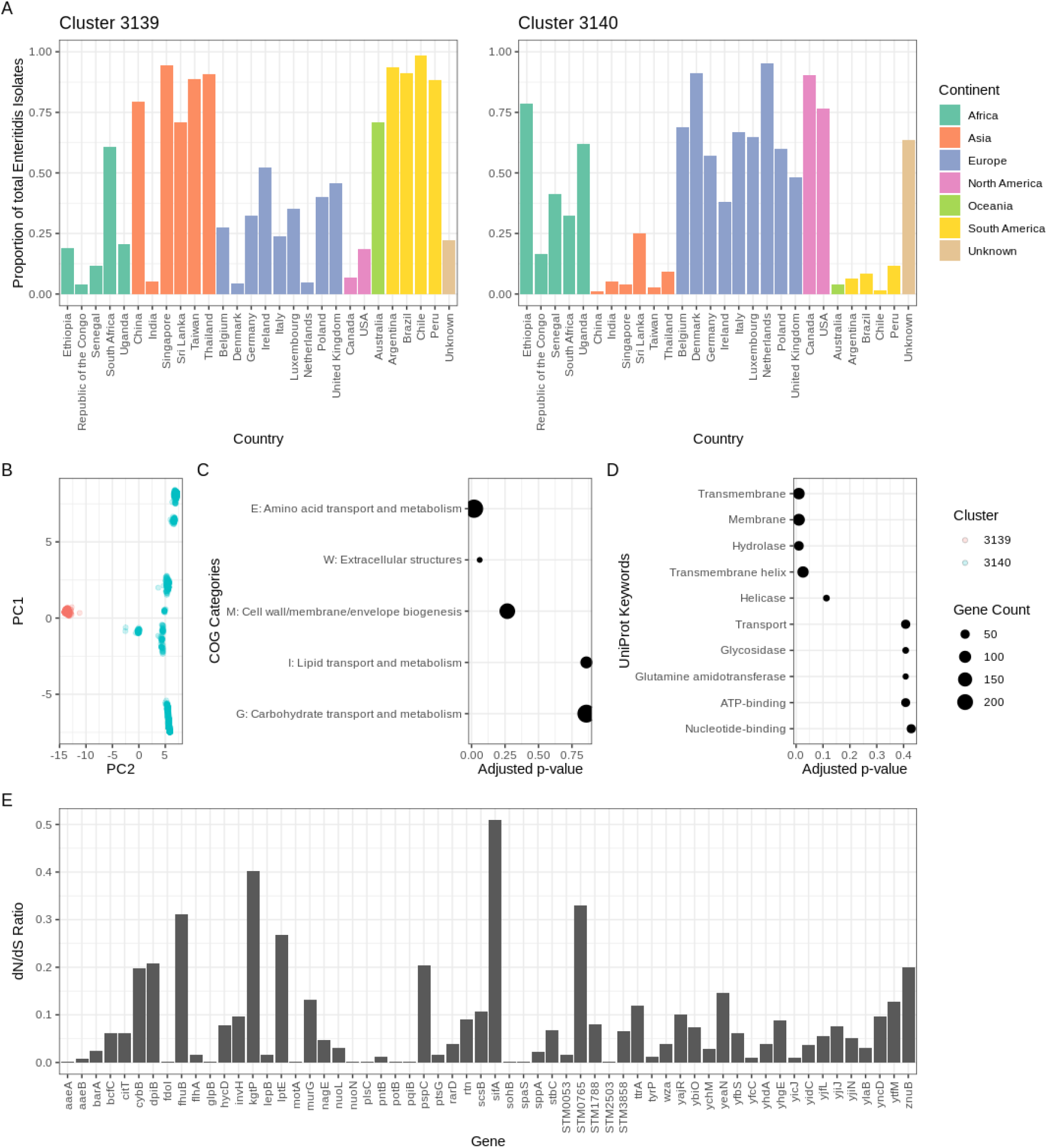
Genomic analysis of geographically segregated Enteritidis subpopulations. A) Geographical distribution of Enteritidis isolates classified in Clusters 3139 and 3140 with the proportion calculated by (isolate count/total Enteritidis isolates count in the dataset). B) PCA analysis of the cgMLST profiles of isolates in the two subpopulations. C and D) Functional annotation enrichment in the top COG categories and UniProt keywords ranked by adjusted p-values. E) dN/dS ratio of the membrane protein genes found to exhibit lineage-specific variation between the two Enteritidis subpopulations.

Subsequent analysis of the principal component loadings identified 284 different alleles involving 168 loci that showed significant contributions (loading threshold > 0.025) to the observed genetic segregation of the two Enteritidis subpopulations. An independent analysis of core genome allelic association to cluster identity using Scoary suggested a higher number of alleles in which 388 alleles involving 183 loci demonstrated significant statistical association (adjusted p-value < 0.05) to Enteritidis subpopulation classification. High concordance was observed between the two methods, as all 168 loci predicted by PCA were also identified in the core genome allelic association analysis. Notably amongst the significant loci, the cpsG gene matched the minimal SNP-based subtyping scheme previously described by Zheng et al. (42) who proposed the differentiation of the two major Enteritidis lineages based on two genes (cpsG and citT) in the *Salmonella* genome (42). Mann-Whitney U-test and DAVID were used to assess functional and pathway enrichment of significant genes which revealed enrichment in COG category E (amino acid transport and metabolism) and membrane protein genes, respectively (Fig. 4C and 4D). Targeted assessment of nonsynonymous (dN) and synonymous (dS) mutation rates of the membrane protein genes revealed a dN/dS ratio < 1 across all membrane protein genes (Fig. 4E). Pan-genome wide association analysis led to the discovery of 6 genes unique to Cluster 3139 with functions in translation, membrane transport, and DNA recombination. (Table S3). Additionally, 6 other genes were found to be unique to Cluster 3140 with functions in DNA recombination, metabolism, and prophage assembly (Table S3).

### 6.5. Comparison of SISTR in-silico serotyping to reported serovars

The increased adoption of WGS for pathogen characterization and the unsustainable costs associated with traditional serology have prompted the replacement of traditional serotyping with in-silico serotyping (43). However, both in-silico serotyping tools such as SISTR (12) or SeqSero2 (44) and the serovar reports in public sequence databases are not flawless due to variable sequencing data quality and human errors, leading to the dissemination of false information and misinterpretations. Misclassification by in-silico serotyping could be a major source of error in this study leading to the overestimation of genetic variability of serovars and hence there is a critical need for a systematic evaluation of in-silico serotyping. According to a previous study that evaluated the performance of in-silico serotyping and the reliability of public database metadata across 42,400 genomes (45), it was reported that SISTR predictions showed 91.91% concordance rate with reported serovars. Moreover, the results discordance was largely attributed to misreports in records and seldom in-silico misclassifications (45).

Here, we took the opportunity to evaluate 102,857 public *Salmonella* genomes archived with serovar information to assess the consistency between SISTR in-silico serotyping and reported serotypes found in public databases. Based on strict string matching, an overall 90% concordance between predicted serovars and reported serovars was observed. We noticed a higher rate of discrepancy in lower quality genome assemblies (< 297/330 hard core genes) in which discrepancies were detected in 32% of lower quality genomes compared to 9.6% in high quality genomes. Further analysis of the discrepancies in high quality genomes revealed sources of inconsistencies unrelated to data quality. 67% of the observed serovar discrepancies in high quality genomes occurred due to the lack of data harmonization for serovar information in public sequence databases. The inconsistent use of different phrases to describe the same serovar and spelling errors were frequently observed. A prominent example is the serovar monophasic variant of Typhimurium which was reported in 8 different ways in the metadata including “I 1,4,[5],12:i:-”, “typhimurium monophasic”, “typhimurium - monophasic”, “4,[5],12:i:-”, “1,4,[5],12:i:-”, “I 4,5,12:i:-”, “4,5,12:i”, and “4,5,12:i-”. However, the remaining 33% of the discrepancies are likely unrelated to data quality or semantics, and instead related to potential errors in in-silico serovar prediction or reports in public data. Examples include the aforementioned O-6 antigen serovar pairs and 1,238 other unique mismatches listed in Table S4. Following the correction for discrepancies due to data quality and semantics, the overall concordance rate between serotyping reports and predictions increased to 96%, suggesting that misclassification by in-silico serotyping has a minor effect on our analyses.

## 7. Discussion

From the core genome allele comparisons of a diverse *Salmonella* dataset of 180,098 genomes, we demonstrated that the global population structure of the pathogen can be arranged into 1,342 stable subpopulations. We confirmed the validity of our population structure analysis based on in-silico serovar classifications in which a crude correlation between serotyping and genomic variation was observed. Moreover, the observation that many serovars subdivided into multiple lineages was consistent with our expectation, considering that the WKL scheme was intended to provide a low resolution typing for *Salmonella* (5). However, the inconsistencies of *Salmonella* serotyping became apparent when the extent of genetic variability in individual serovars was evaluated. Collectively, we highlighted two major types of serotyping inconsistencies: 1) segregation of closely related strains into unrelated antigenic groups and 2) inter-serovar genetic distance exceeding that of intra-serovar comparisons. Importantly, our observations suggested that the inconsistent serovars do not represent groupings of genetically related isolates and thus their corresponding isolates likely do not share the same epidemiological or clinical implications. Our findings are concordant with a previous report that described the failure of the serovar Senftenberg to represent a classification of genetically related isolates (46). Specifically, different subsets of the serovar Senftenberg isolates were reported to share either the same sequence type (ST14) with the serovar Westhampton, or the same sequence type (ST185) with another serovar, Dessau (46). For future work, we suggest a systematic analysis of serotyping inconsistencies in a phylogenetic context to confirm the non-monophyletic nature of the serovars that we have identified solely based on genetic distances.

Indeed although a correlation exists between antigenic and genetic variations, caution must be made surrounding specific serovars to avoid misinterpretations of genetic, biological and epidemiological features based on serotyping. It can be postulated that the occasional discrepancy between antigenic and genome-wide variation is attributable to the homologous recombination of genes involved in antigen synthesis which confounds the inference of genealogy. Recently, it has been reported that the O-antigen biosynthesis genes are situated in a genomic island with distinctive GC content rendering it a hot spot for homologous recombination and horizontal gene transfer (47). The discovery of over 200 serovars that had not been reported as putative non-monophyletic classifications is further indicative of the negligence of genomic investigations for many serovars thus far. Moreover, previous studies often selected a minor subset of genomes to represent a given serovar leading to insufficient coverage of genetic diversity and consequently the assumption of monophyly (48, 49). Hence, the collective analysis of a highly genetically diverse dataset on the scale of hundreds of thousands of genomes remains valuable to deduce a more holistic view of bacterial population structures.

To contribute to the knowledge base of *Salmonella* serovar classification, we compiled a public resource (Table S5) that documents the putative non-monophyletic serovars, as well as the serovars that belong to the same lineage as the putative non-monophyletic serovars. The deficiencies of serotyping highlighted in this study can be addressed by WGS given its capability to survey genome-wide markers and thereby empower genomics researchers to model the divergence of bacterial populations with greater precision and resolution. By subdividing *Salmonella* into 1,342 groups based on the most stable *T,* a genomic-based typing scheme can be developed. A single sequence can be selected from each stable cluster to construct a set of reference sequences that would collectively represent the global *Salmonella* population structure. Subsequent alignments of query sequences to the reference sequences would thus inform the subtypes of the query genomes based on the most similar reference.

In fact, a similar solution has already been implemented in Enterobase that integrates the application of cgMLST thresholds to define stable *Salmonella* clusters represented by hierarchical cluster codes (16). The Enterobase cgMLST thresholds have been specifically chosen with the intention to reflect natural *Salmonella* subpopulations such as subspecies, superlineages, and eBGs (16). The integration of hierarchical cluster code assignments in Enterobase represents an initial step towards the establishment of a standardized WGS-based typing scheme for *Salmonella*. However, a global community consensus on a standardized genomic-based typing scheme remains critical to encourage the usage of a consistent language to share *Salmonella* population dynamics across jurisdictions and international borders. Moreover, community consensus of a core or whole genome MLST scheme is also necessary to converge on a fixed set of genome-wide loci for comparison and reinforce a standardized allele number nomenclature. In particular, the Chewie Nomenclature Server (https://chewbbaca.online/) could be a highly promising avenue to synchronize local schema with globally maintained schema and centralize a collection of gene-by-gene subtyping schema for microbial pathogens (50).

The importance of standardization and data harmonization cannot be undermined, as the lack of controlled vocabularies in open data settings hinders data interpretability and summarization. The high degree of metadata inconsistencies in public foodborne pathogen databases is evident from the frequent discrepancies between SISTR predictions and reported serovars solely due to semantics. The unrestrictive entry of serovar information in GenomeTrakr and PubMLST hinders an accurate estimation of the number of serovars in the study dataset based on the reported serovars in the metadata alone. Moreover, it renders the estimated 90% concordance rate between SISTR predictions and reported serovars to likely represent an underestimation of the true concordance rate. The integration of controlled vocabularies for subtype information in public databases would be highly valuable for future studies that intend to analyze public *Salmonella* sequences by eliminating the necessity for time-consuming data harmonization and thus reinforcing analysis reproducibility.

To demonstrate the additional utility of nAWC to reveal cluster consolidation dynamics and inform evolutionary histories, we analyzed the genomic differences between two geographically segregated Enteritidis lineages that converged at a cluster destabilizing threshold (*T* = 157). The pronounced allele frequency bias across different geographical regions demonstrated from genome-wide association analyses suggested that the observed lineage divergence could be a consequence of niche adaptation or founder effects. A previous study on the evolutionary history of *Salmonella* Enteritidis had estimated the most recent common ancestor of present-day Enteritidis lineages to have existed in the 1600s when cross-continental overseas trades were increasingly embraced (51). Hence, an ancestral European Enteritidis subpopulation may possibly have migrated to America during the era of colonial trade and evolved in reproductive isolation leading to the fixation of the lineage-specific alleles observed in Cluster 3139. However, the enrichment of allelic bias in metabolic and membrane transport functions could reflect the local adaptation of the Enteritidis subpopulations to the variabilities in regional nutrient availability, a phenomenon that has been previously reported in a metabolic analysis of Enteritidis lineages in high- and low-income settings (52). We recommend a future analysis of in-vitro metabolic activities of the two Enteritidis populations and correlating differential metabolic activities to the divergence of lineage-specific alleles to test our hypotheses.

In the genomics era, the development of efficient bioinformatics tools and high-throughput sequencing have revolutionized our understanding about microbial evolution and biology. Here, we continued to echo the value of genomic data by proposing novel ways to define the global *Salmonella* population structure and discovering novel genetic patterns that differentiate microbial subpopulations. Our study is a further demonstration of how global genomic data sharing benefits scientific research. The open accessibility of large-scale genomic datasets with minimal metadata fields will foster new directions to investigate molecular evolution and microbial diversity. With increasing integration of genomics in infectious disease surveillance worldwide, we encourage joint efforts across scientific research and public health to standardize a genomic-based typing nomenclature for global disease reporting such that *Salmonella* classifications consistently reflect both genetic relatedness and epidemiological significance.

## 8. Author Statements

### 8.1. Authors and contributors

C.L.: Conceptualization, formal analysis, methodology, validation, software, visualization, funding acquisition, and writing – original draft W.H.: Project administration, conceptualization, supervision, funding acquisition, and writing – review and editing

## 8.4. Acknowledgements

The authors thank Zohaib Anwar for his constructive feedback on the original manuscript and the SISTR developers: John Nash, Ed Taboada, and James Robertson for their insightful discussions on the subject matter. The authors express gratitude to all the contributors to GenomeTrakr and PubMLST whose data enabled this large scale analysis.

## 9. Figure Legends

**Figure S1.**
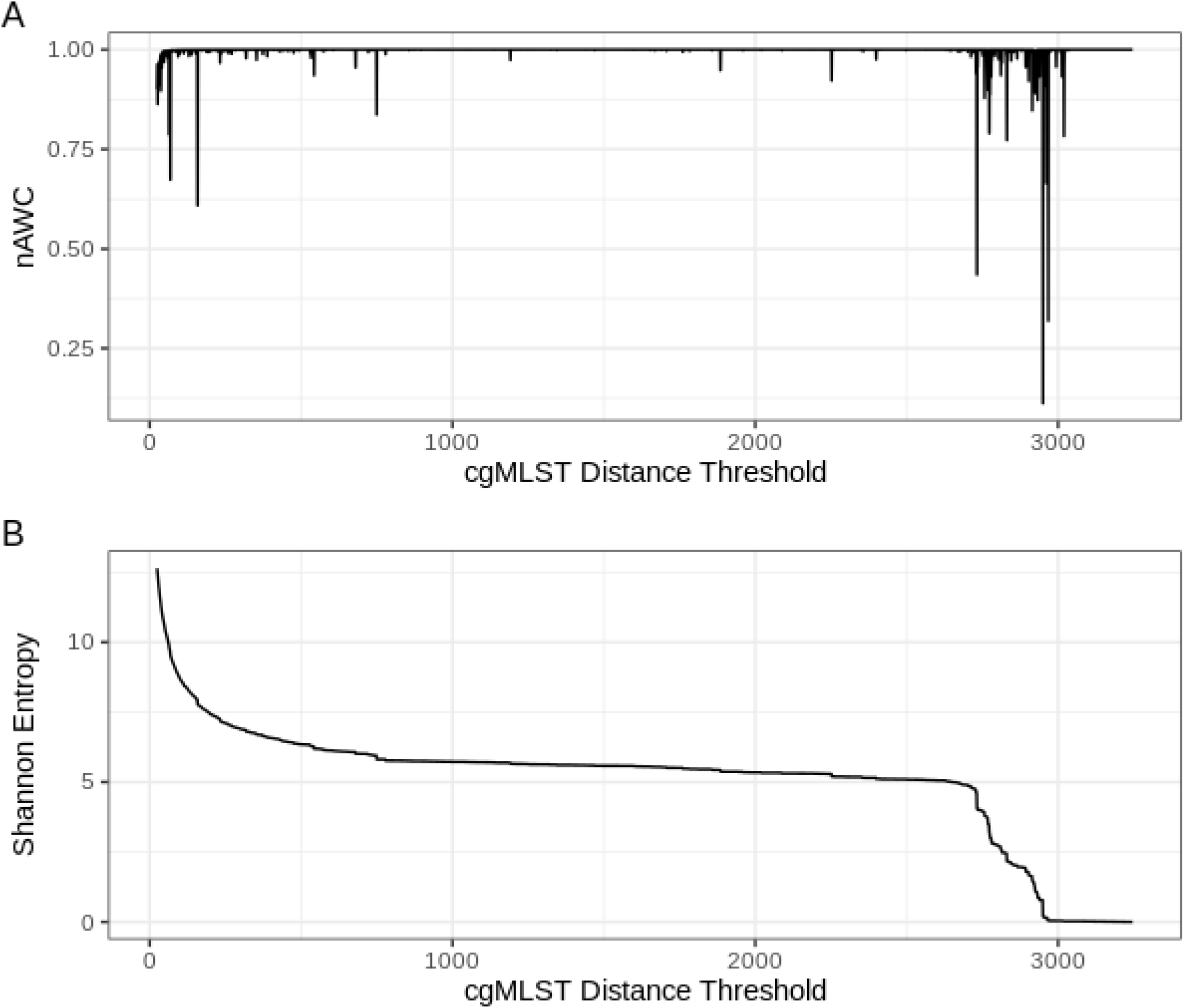
nAWC (A) and Shannon entropy (B) plots calculated across the range of all possible cgMLST distance thresholds.

**Figure S2.**
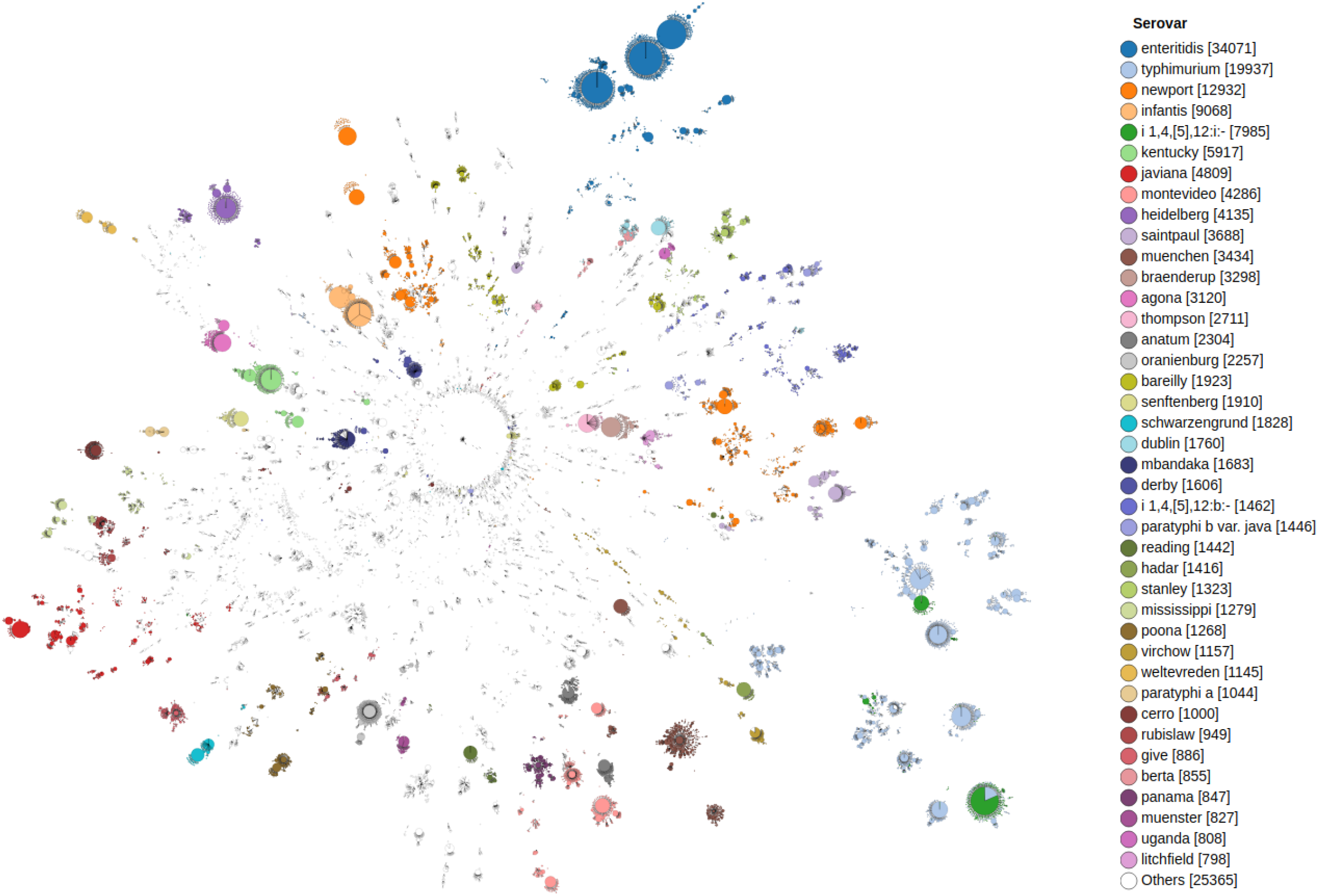
Unrooted neighbour-joining tree constructed from 3,246-loci cgMLST profiles of 180,098 *Salmonella enterica* genomes with color annotations reflecting serovar classifications only. The tree is visualized using GrapeTree (43)

**Figure S3.**
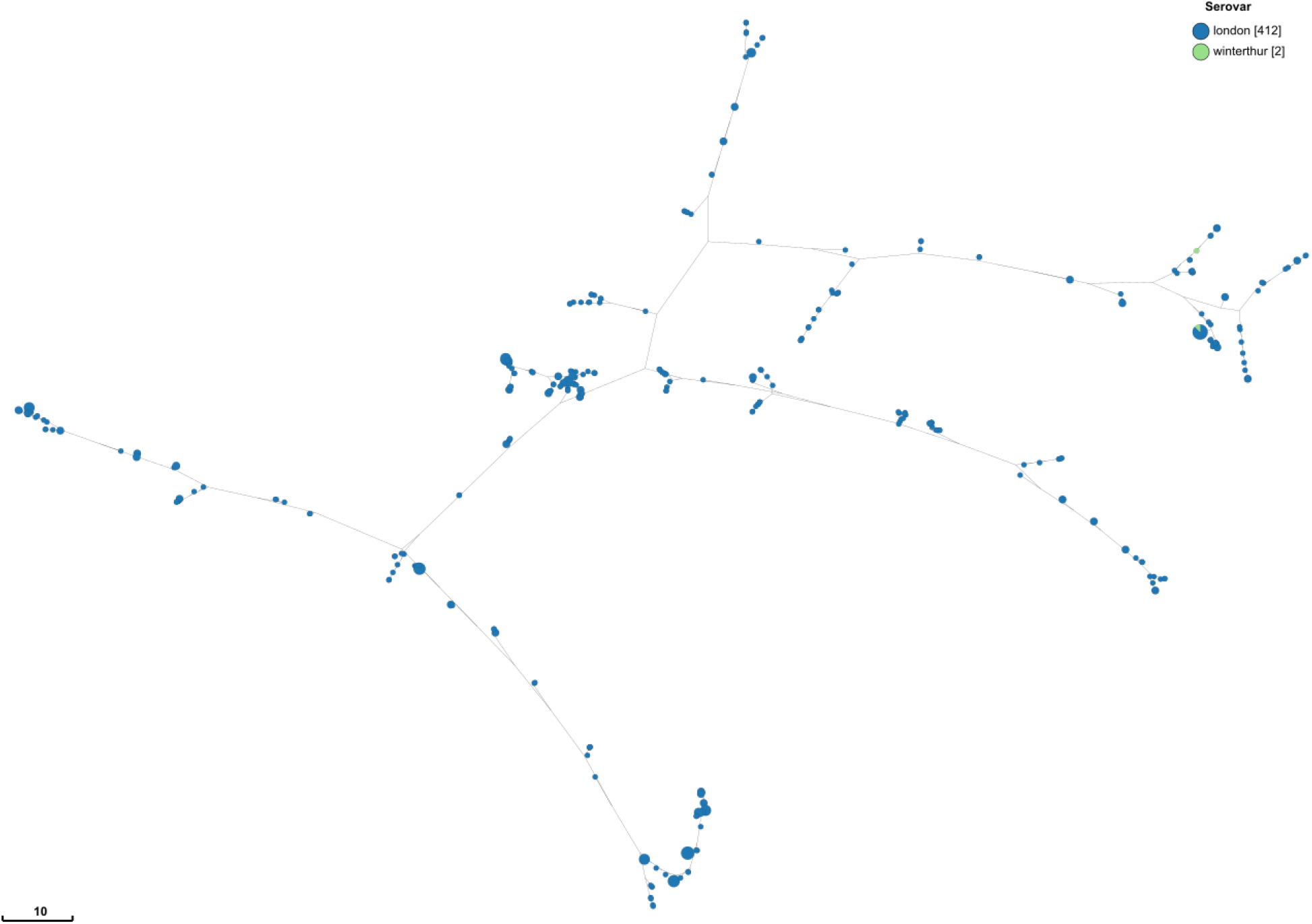
Unrooted maximum likelihood tree of serovars London and Winterthur genomes visualized in GrapeTree. The branch lengths in the visualization are not to scale of the actual lengths.

**Figure S4.**
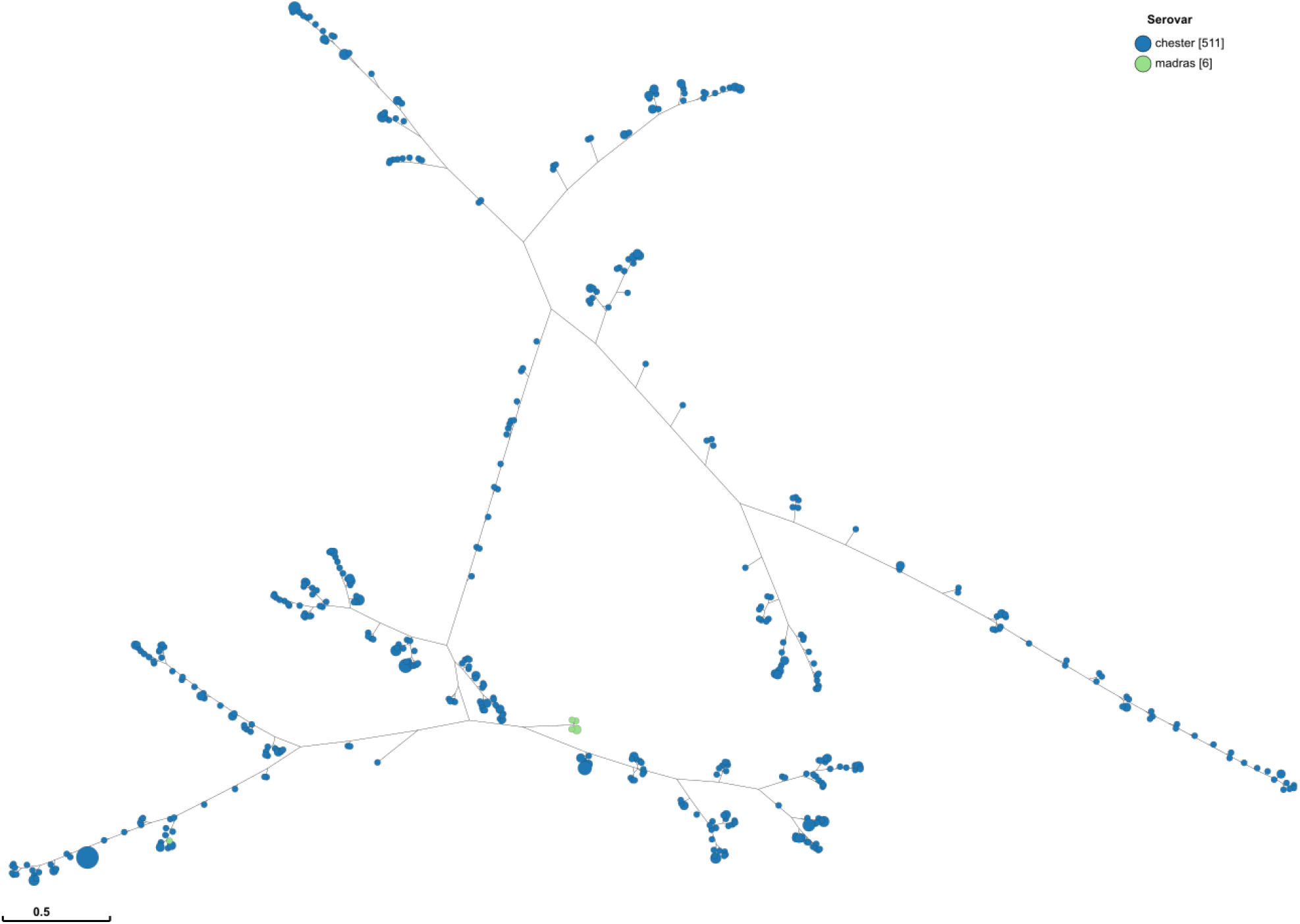
Unrooted maximum likelihood tree of serovars Chester and Madras genomes visualized in GrapeTree. The branch lengths in the visualization are not to scale of the actual lengths.

**Figure S5.**
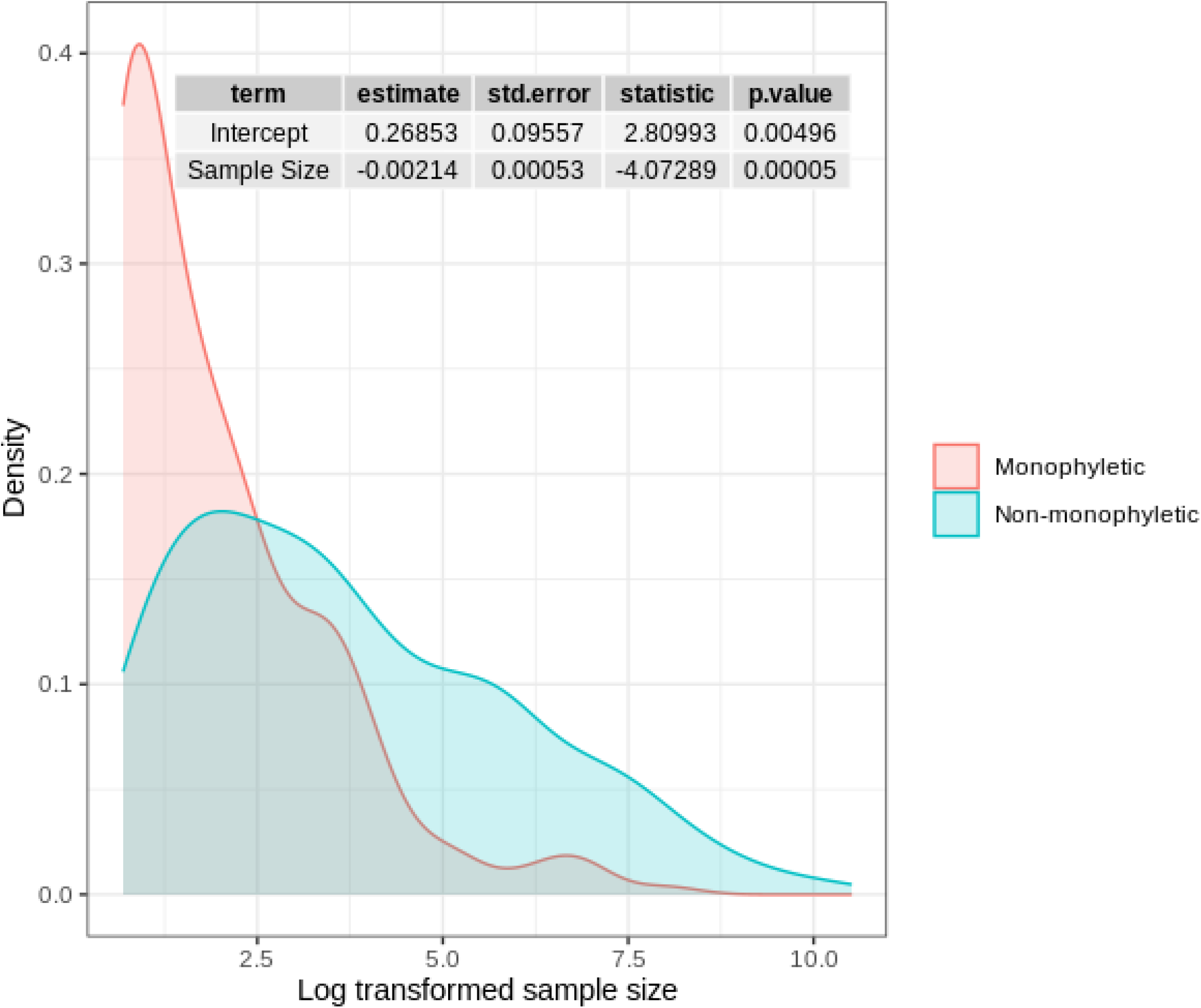
Sample size density distribution of predicted monophyletic and non-monophyletic serovars. The summary table shows various statistics including the statistical significance and regression coefficient of serovar sample size as a predictor of monophyly based on logistic regression.

